# Role of the drug efflux pump in the intrinsic cefiderocol resistance of *Pseudomonas aeruginosa*

**DOI:** 10.1101/2022.05.31.494263

**Authors:** Sota Ikawa, Seiji Yamasaki, Yuji Morita, Kunihiko Nishino

## Abstract

Cefiderocol is a novel siderophore cephalosporin antibiotic exhibiting activities against carbapenem-resistant Gram-negative bacteria including *Pseudomonas aeruginosa* and *Enterobacteriaceae*. Drug efflux pumps are reportedly involved in both intrinsic and acquired drug resistance, although their role in bacterial cefiderocol susceptibility remains poorly understood. In this study, we investigated how drug efflux pumps contribute to bacterial cefiderocol susceptibility using the efflux pump(s)-deficient and overexpressing strains of *P. aeruginosa, Escherichia coli*, and *Salmonella enterica*. We observed that the *mexAB*-*oprM*-deficient *P. aeruginosa* mutant displayed increased cefiderocol susceptibility compared to the wild-type strain. The overexpression of *mexAB-oprM* or *mexXY-oprM* in the *mexAB*-*oprM*-deficient mutant increased the MIC value of cefiderocol. Furthermore, the pump inhibitor phenylalanine-arginine β-naphthylamide increased cefiderocol susceptibility in wild-type *P. aeruginosa* whereas it did not affect the susceptibility of the *mexAB*-*oprM*-deficient mutant. These data indicate that the MexAB-OprM drug efflux system contributes to the intrinsic cefiderocol resistance of *P. aeruginosa*. In addition, MexXY-OprM partially complemented the function of MexAB-OprM in the cefiderocol susceptibility, when expressed.

## Introduction

The emergence of drug-resistant bacteria is a current global problem. WHO has published a list of drug-resistant organisms that represent a serious threat to human health, and called for the development of new antimicrobial agents against them. *Acinetobacter baumannii, Pseudomonas aeruginosa*, and *Enterobacteriaceae* have been listed in the highest priority category as organisms that acquired resistance against various antimicrobial agents, including carbapenems and third-generation cephalosporins (1). The resistance mechanisms against carbapenems include hydrolysis of antimicrobials by carbapenemases, reduced porin expression or mutation of porins, and efflux pump overexpression (2).

Cefiderocol is a novel siderophore cephalosporin antibacterial agent that effectively crosses the outer membrane of Gram-negative bacteria, including that of multidrug-resistant bacteria (3). Cefiderocol is the only drug so far that exhibits antimicrobial activity against carbapenem-resistant *A. baumannii, P. aeruginosa*, and *Enterobacteriaceae* (4). In addition, cefiderocol, as a siderophore, possesses an iron-binding structure that allows its active bacterial uptake through iron uptake mechanisms, and it could effectively inhibit cell wall synthesis in the periplasm. Cefiderocol was suggested to be unaffected by the three carbapenem resistance acquisition-associated mechanisms: mutated porin channel-related reduced membrane permeability, β-lactamase-mediated inactivation, and efflux pump overproduction (5, 6). Previous studies examined the antibacterial activity of cefiderocol against various drug-resistant bacteria and showed its higher potency against β-lactamase-possessing Gram-negative bacteria compared to carbapenems, including ESBL and carbapenemases. The presence or absence of porins, related to carbapenem susceptibility, has also been suggested to leave cefiderocol activity unaffected (7). Moreover, the iron transporters reportedly affect cefiderocol activity (3). However, the role of drug efflux pumps in bacterial cefiderocol susceptibility remains elusive.

Previous studies described the presence of numerous drug efflux pumps in Gram-negative bacteria including *E. coli, Salmonella*, and *P. aeruginosa*. Among them, the drug efflux system of the resistance-nodulation division (RND) superfamily plays a particularly important role, such as AcrAB-TolC in *Salmonella* and *E. coli* or MexAB-OprM in *P. aeruginosa*, recognizing and exporting a wide variety of structurally unrelated antimicrobial agents (8–11).

In this study, we first measured and compared the cefiderocol MIC values of wild-type *E. coli, S. enterica*, and *P. aeruginosa* strains and multiple drug efflux pump-deleted mutants using iron-limited media. Moreover, we examined the changes in susceptibility when *mexAB-oprM* or *mexXY-oprM* was expressed in *P. aeruginosa mexAB-oprM*-deleted mutants. Furthermore, to investigate how pump inhibition affects the cefiderocol susceptibility of *P. aeruginosa*, we used phenylalanine-arginine β-naphthylamide to monitor the changes in susceptibility. Based on the obtained results, our study clarifies the role of the drug efflux pump in the intrinsic cefiderocol resistance of *P. aeruginosa*.

## Results

### Effect of efflux pump deletions on the cefiderocol susceptibility of *E. coli, S. enterica*, and *P. aeruginosa*

In order to investigate the contribution of drug efflux pumps to cefiderocol susceptibility, we measured cefiderocol MIC values against the wild-type and efflux pump-deleted mutants of *E. coli, S. enterica*, and *P. aeruginosa* (Table 1). We measured the MIC values both in cation-adjusted and iron-depleted cation-adjusted Muller-Hinton broths ((CAMHB and ID-CAMHB, respectively). We detected no significant difference in the cefiderocol susceptibility between the wild-type *E. coli* strain and the mutant lacking five RND efflux systems (Table 1). Furthermore, we observed no difference either in the susceptibility of the wild-type *S. enterica* strain and that of the mutant lacking nine efflux pumps. The number of drug efflux pumps, including AcrAB, AcrD, MdtABC, MdtEF, AcrEF, MdsAB, EmrAB, and MacAB, can use the TolC outer membrane protein for drug efflux in both *E. coli* and *S. enterica* (12,13). We also measured the cefiderocol susceptibility of the *tolC*-deleted *E. coli* and *S. enterica* mutants. However, the MIC values against these mutants were the same as in the case of the wild-type strains. In contrast to the results obtained in *E. coli* and *S. enterica*, we observed a difference in the cefiderocol susceptibility between the wild-type *P. aeruginosa* and the efflux pump-deleted mutants. For example, the mutant lacking five drug efflux systems, including *mexAB-oprM, mexXY, mexCD-oprJ, mexEF-oprN*, and *mexHI-opmD*, was 32-fold more susceptible to cefiderocol than the wild-type in ID-CAMHB. The cefiderocol MIC values against the *P. aeruginosa* strains in CAMHB were higher than those in ID-CAMHB. Furthermore, smaller differences could be observed between the wild-type and the efflux pump-deleted strains. This was probably due to the good cefiderocol uptake by *P. aeruginosa* under iron limitation. In such a situation, the presence or absence of a drug efflux pump might have affected more the drug accumulation, observed as a difference in susceptibility. The *mexAB* or *mexAB-oprM* genes were deleted in all the mutants susceptible to cefiderocol, suggesting that the MexAB-OprM efflux system is involved in the intrinsic cefiderocol resistance under iron-limited conditions. In fact, the *mexAB-oprM*-deleted mutant was 16-fold more susceptible to cefiderocol than the wild-type strain. *P. aeruginosa* growth in ID-CAMHB was slower than in CAMHB, although no significant difference could be detected between the wild-type strain and the *mexAB-oprM*-deleted mutant under both conditions (Fig. 1), suggesting that the difference in cefiderocol susceptibility between them is probably due to the MexAB-OprM-mediated cefiderocol efflux.

**Table 1.** Effects of the drug efflux pump deletion on cefiderocol susceptibility

| Organism | MIC (mg/L) |  |
| --- | --- | --- |
|  | Cefiderocol |  |
|  | CAMHB | ID-CAMHB |
| *E.coli* |  |  |
| Wild-type | 0.5 | 0.25 |
| $\Delta tolC$ | 0.25 | 0.25 |
| $\Delta acrB\ acrC\ mdtABC\ mdtEF\ acrEF$ | 0.25 | 0.25 |
| *Salmonella* |  |  |
| Wild-type | 0.25 | 0.25 |
| $\Delta tolC$ | 0.25 | 0.25 |
| $\Delta acrAB\ acrEF\ acrD\ mdtABC\ mdsABC\ emrAB\ mdfA\ mdtK\ macAB$ | 0.12 | 0.25 |
| *P. aeruginosa* |  |  |
| Wild-type | 1 | 0.5 |
| $\Delta mexXY$ | 1 | 0.5 |
| *ΔmexAB* | 0.5 | 0.062 |
| *ΔmexAB-oprM* | 0.5 | 0.03 |
| *ΔmexCD-oprJ* | 1 | 0.5 |
| *ΔmexXY mexAB* | 0.5 | 0.062 |
| *ΔmexXY mexAB-oprM* | 0.5 | 0.062 |
| *ΔmexXY mexCD-oprJ mexEF-oprN* | 2 | 0.5 |
| *ΔmexAB mexCD-oprJ mexEF-oprN* | 0.5 | 0.062 |
| *ΔmexAB-oprM mexXY mexCD-oprJ* | 1 | 0.016 |
| *ΔmexAB-oprM mexXY mexEF-oprN* | 1 | 0.008 |
| *ΔmexAB mexXY mexCD-oprJ mexEF-oprN* | 1 | 0.03 |
| *ΔmexAB-oprM mexXY mexCD-oprJ mexEF-oprN* | 1 | 0.03 |
| *ΔmexAB-oprM mexXY mexCD-oprJ mexEF-oprN mexHI-opmD* | 0.5 | 0.016 |

**Figure 1.**
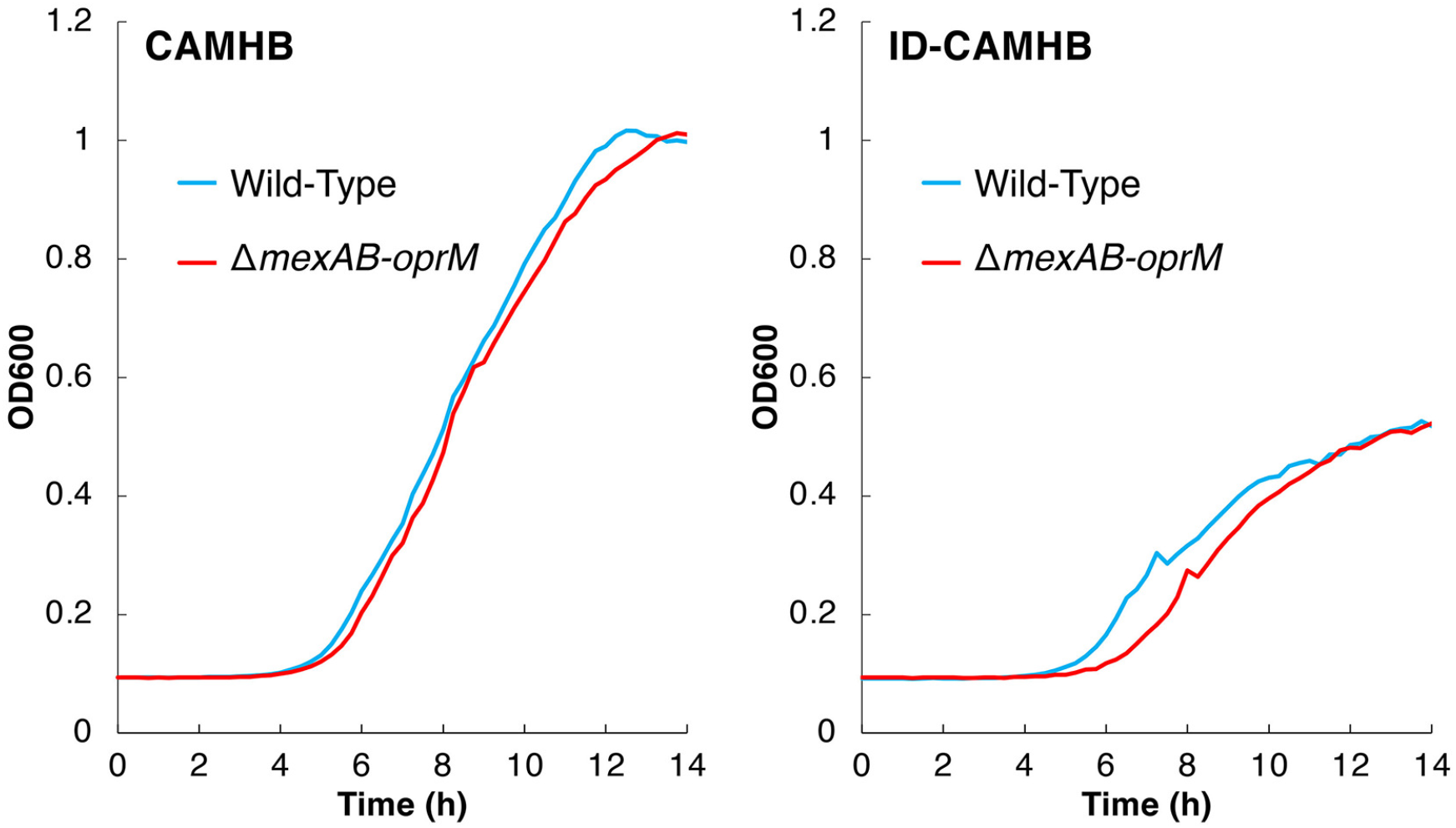
Growth of *P. aeruginosa* PAO1 wild-type and Δ*mexAB-oprM* strains in cation-adjusted Mueller-Hinton broth (CAMHB) (left column) or iron-depleted cation-adjusted Mueller-Hinton broth (ID-CAMHB) (right column). The growth of PAO1 and YM5 was measured as described in the Materials and Methods.

### Effect of MexAB-OprM and MexXY-OprM expression on the cefiderocol susceptibility of *P. aeruginosa*

As described above, the *mexAB-oprM*-deleted *P. aeruginosa* mutant was more susceptible to cefiderocol than the wild-type strain. To see the potential complementary effect of the *mexAB-oprM* expression, we transformed a plasmid carrying *mexAB-oprM* into the *mexAB-oprM*-deleted mutant and measured MIC. The transformation with the *mexAB*-*oprM* expression plasmid restored cefiderocol susceptibility to the level of the wild-type strain (Table 2). Similarly, the deletion of *mexAB*-*oprM* also reduced the MICs of antimicrobial agents other than imipenem, and they got restored upon the *mexAB*-*oprM* expression plasmid transformation (Table 2). Regarding imipenem, we observed no change in MIC in the presence or absence of *mexAB-oprM* (Table 2), consistently with a previous study reporting that imipenem was not affected by the efflux pump but the OprD expression level (14).

**Table 2.** Effect of *mexAB-oprM* or *mexXY-oprM* on *P. aeruginosa* drug susceptibility

| Organism | MIC (mg/L) |  |  |  |  |  |  |  |
| --- | --- | --- | --- | --- | --- | --- | --- | --- |
|  | Cefiderocol |  | Ceftazidime | Cefepime | Meropenem | Imipenem | Norfloxacin | Chloramphenicol |
|  | CAMHB | ID-CAMHB |  |  |  |  |  |  |
| *P. aeruginosa* |  |  |  |  |  |  |  |  |
| Wild-type | 1 | 0.5 | 2 | 2 | 0.5 | 1 | 0.5 | 64 |
| *ΔmexAB* | 0.5 | 0.062 | 1 | 1 | 0.12 | 1 | 0.25 | 16 |
| *ΔmexAB-oprM* | 0.5 | 0.03 | 1 | 0.25 | 0.12 | 1 | 0.062 | 4 |
| *ΔmexAB-oprM*/vector(pMMB67HE) | 0.5 | 0.03 | 1 | 0.5 | 0.12 | 1 | 0.12 | 4 |
| *ΔmexAB-oprM*/pmexAB-*oprM* | 1 | 0.5 | 2 | 2 | 0.5 | 1 | 0.5 | 128 |
| *ΔmexAB-oprM*/pmexXY-*oprM* | 1 | 0.12 | 1 | 8 | 0.25 | 1 | 2 | 16 |

As MexXY is also known to form a functional complex with OprM (15) and it is also an important efflux system expressed in multidrug-resistant *P. aeruginosa* strains (16,17), we also investigated the complementary effect of *mexXY-oprM* expression. When we transformed the plasmid carrying *mexXY-oprM* into the Δ*mexAB-oprM* mutant, we observed a four-fold increase in the cefiderocol MIC value, similar to ceftazidime, meropenem, and chloramphenicol (Table 3). Therefore, we could conclude that MexXY-OprM is involved in exporting cefiderocol, meropenem, ceftazidime, and chloramphenicol in a manner that partially complements the function of MexAB-OprM. In addition, *mexXY-oprM* expression increased cefepime and norfloxacin resistance beyond the wild-type level. Therefore, MexXY-OprM is considered to be the efflux system that can efficiently recognize and export cefepime and norfloxacin.

**Table 3.**
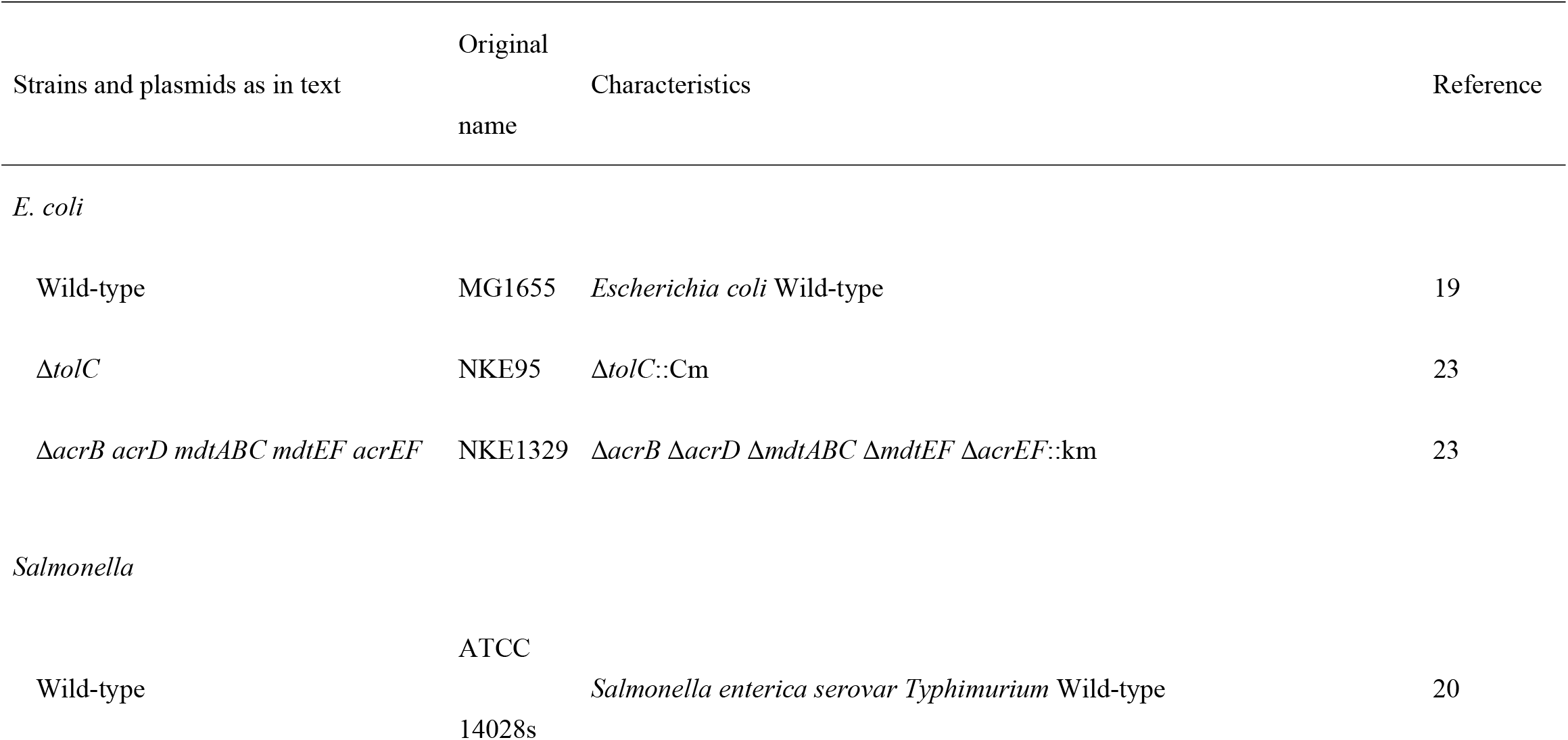
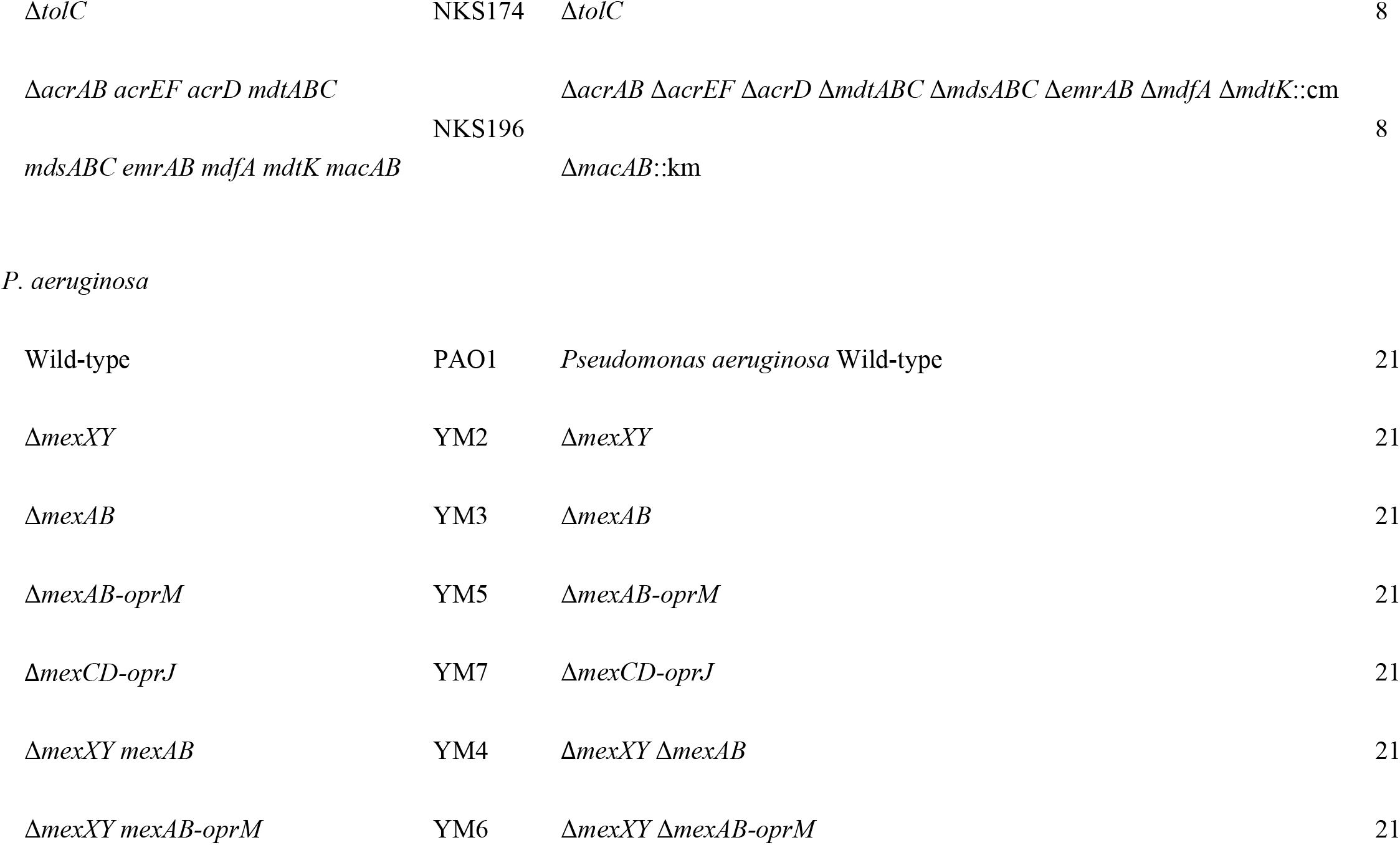
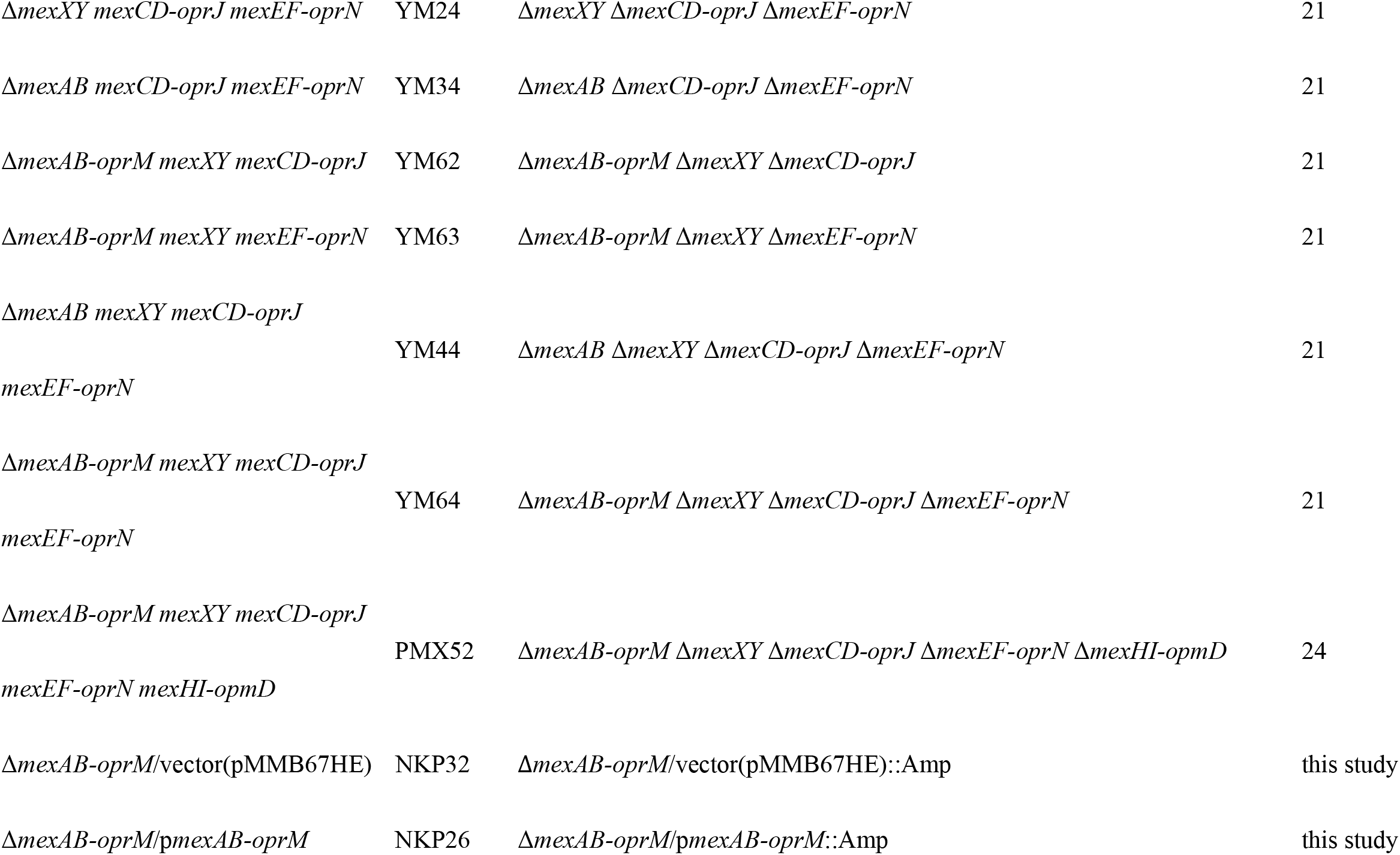
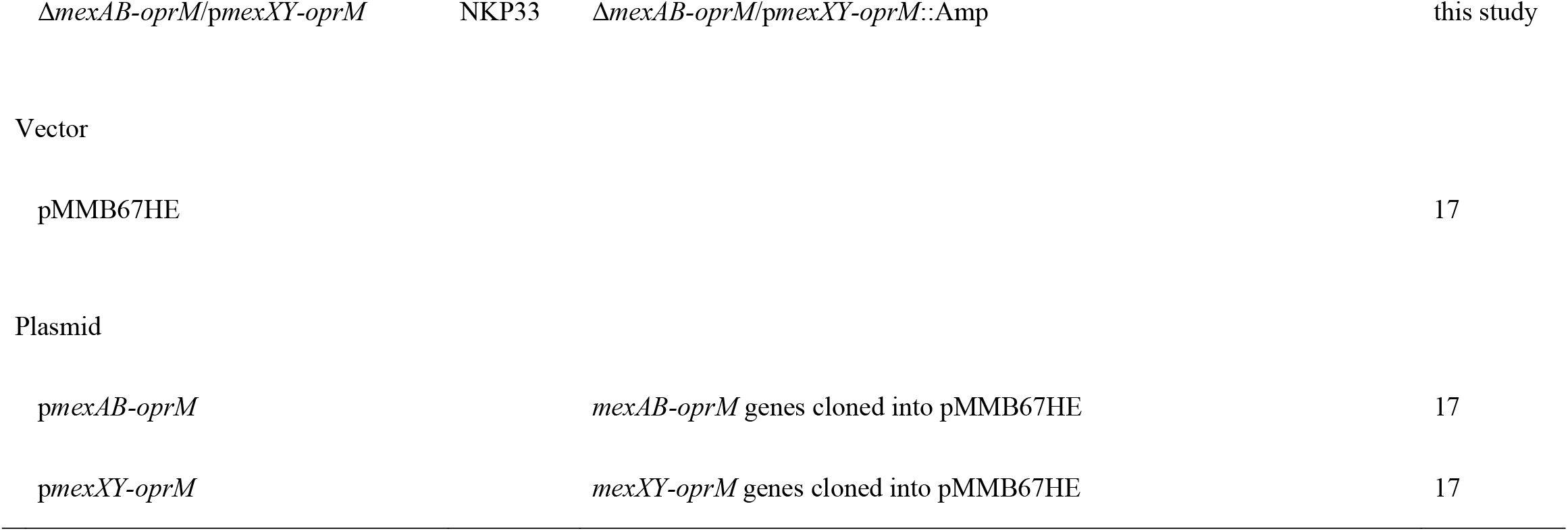
Bacterial strains and plasmids used in this study.

### Effect of the efflux pump inhibitor on the cefiderocol susceptibility of *P. aeruginosa*

The aforementioned results indicate that MexAB-OprM is involved in the intrinsic cefiderocol resistance in *P. aeruginosa*, suggesting that cefiderocol might exhibit its bactericidal function more effectively against *P. aeruginosa* upon functional MexAB-OprM inhibition. We thus evaluated the synergistic effects of the efflux pump inhibitor phenylalanine-arginine *β*-naphthylamide (PA*β*N) and cefiderocol against *P. aeruginosa* using the checkerboard method (Fig. 2). In the wild-type strain, we observed a PA*β*N concentration-dependent increase in cefiderocol susceptibility with a 16-fold increase in the presence of 32 mg/L of PA*β*N (Fig. 2). However, we observed no change in cefiderocol sensitivity with Δ*mexAB-oprM* in the presence of PA*β*N (Fig. 2). These results indicated that the intrinsic cefiderocol resistance of *P. aeruginosa* could be suppressed by inhibiting the function of the drug efflux pump.

**Figure 2.**
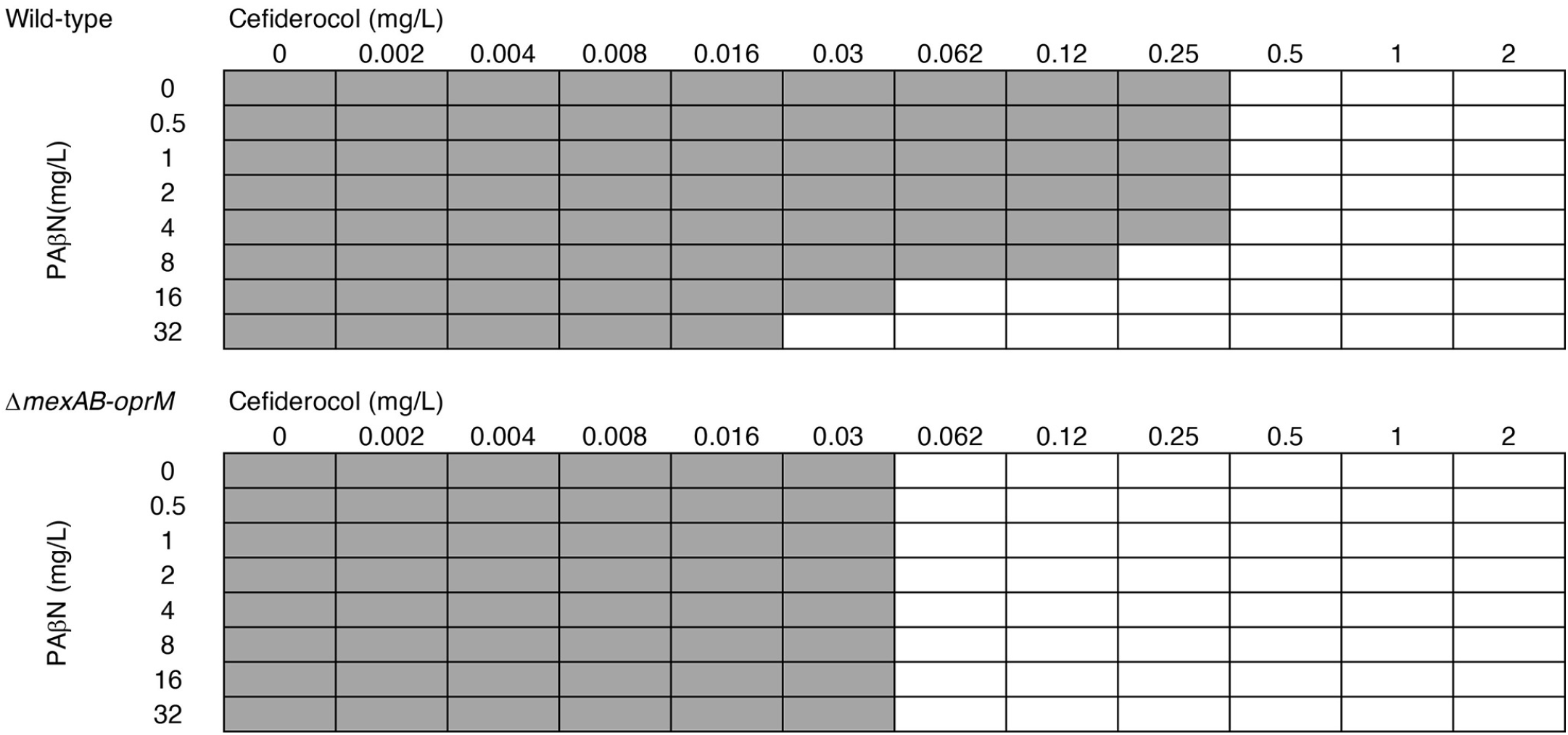
Synergistic effect of cefiderocol and the efflux inhibitor PA*β*N against *P. aeruginosa*. To evaluate the effect of the combination of cefiderocol and PA*β*N against PAO1 (wild-type) and Δ*mexAB-oprM* (YM5), we performed the checkerboard assay as described in the Materials and Methods.

## Discussion

This study demonstrates the involvement of the MexAB-OprM drug efflux system in the intrinsic cefiderocol resistance of *Pseudomonas aeruginosa*. MexXY-OprM is also reportedly involved in cefiderocol efflux in a manner that partially complements the function of MexAB-OprM. Probably because MexAB-OprM masks MexXY function or the *mexXY* expression level is low in the wild-type strain, the *mexXY* deletion itself did not alter cefiderocol susceptibility. As MexXY is reportedly induced in the presence of certain drugs (18), if a MexXY-OprM is expressed in a strain in which MexAB-OprM is no longer functional, this efflux system is suggested to be also involved in intrinsic cefiderocol resistance.

As for the MexCD-OprJ and MexEF-OprN efflux systems, we observed no change in the cefiderocol susceptibility of *P. aeruginosa* when these genes were deleted from the *mexXY*-deficient strain, suggesting that neither of these efflux systems is involved in intrinsic cefiderocol resistance in the PAO1 strain. In *E. coli* and *S. enterica*, no changes could be detected in cefiderocol susceptibility between the wild-type and multiple drug efflux gene-lacking strains, suggesting that the drug efflux systems of these two bacterial species are not related to the intrinsic cefiderocol resistance. MexAB-OprM plays a significant role in the intrinsic resistance of *P. aeruginosa*, probably due to its high cefiderocol efflux activity.

The chemical structure of cefiderocol is similar to both ceftazidime and cefepime, the major difference is the addition of a chlorocatechol group on the end of the C-3 side chain, conferring the siderophore activity (7). When we compared ceftazidime, cefepime, and cefiderocol for their MIC reduction in the lack of *mexAB*-*oprM*, this reduction was higher for cefiderocol (ceftazidime and cefepime: 2-fold decrease, cefiderocol: 16-fold). Therefore, the unique siderophore structure of cefiderocol could facilitate its recognition by MexAB-OprM.

Previous studies have shown that mutations in *mexR* and *nalD*, resulting in the increased expression of *mexAB*-*oprM*, led to a 2-fold increase in the cefiderocol MIC value. This is a less potent MIC increase than that related to other agents such as ceftazidime, aztreonam, and ciprofloxacin, the authors thus concluded the lack of significant impact on the cefiderocol susceptibility along with MexAB-OprM (6). Our study revealed that under the iron-limiting conditions, the loss of MexAB-OprM significantly affected the cefiderocol susceptibility of *P. aeruginosa*, indicating the importance of MexAB-OprM in the intrinsic cefiderocol resistance of this organism.

In this study, we showed that the intrinsic cefiderocol resistance of *P. aeruginosa* can be suppressed by the combination with the efflux inhibitor, suggesting that the inhibition of the MexAB-OprM drug efflux pump can suppress the intrinsic cefiderocol resistance of *P. aeruginosa*. If efflux inhibitors applicable in clinical practice would be developed, a more effective cefiderocol treatment might be possible when used in combination. Finally, it is also possible that the drug efflux overexpression-related future emergence of cefiderocol-resistant strains could be effectively treated in combination with efflux inhibitors.

## Materials and Methods

### Bacterial strains, plasmids, and growth conditions

Table 3 contains all the bacterial strains and plasmids that we used in this study. The *E. coli, S. enterica* serovar Typhimurium, and *P. aeruginosa* strains used in this study were derived from the wild-type strains MG1655 (19), ATCC14028s (20), and PAO1 (21), respectively. The bacterial strains were grown at 37°C in Lysogeny broth, CAMHB, or iron-depleted cation-adjusted Mueller-Hinton broth (ID-CAMHB). Ampicillin was added to the growth media at a final concentration of 100 mg/L for plasmid maintenance. The plasmids pMMB67HE, p*mexAB*-*oprM*, and p*mexXY-oprM* were kindly provided by Dr. Taiji Nakae (17).

### Construction of gene deletion mutants

To construct the gene deletion mutants of *E. coli* and *S. enterica*, gene disruption was performed as described by Datsenko and Wanner (8,22,23). The chloramphenicol resistance *cat* gene and the kanamycin resistance *aph* gene, flanked by Flp recognition sites, were PCR-amplified, and the products were transformed into the recipient MG1655 or ATCC14028s strain harboring plasmid pKD46, expressing the Red recombinase. The mutated loci were PCR-verified and *cat* and *aph* were eliminated using plasmid pCP20. *P. aeruginosa* PAO1-derived strains lacking genes encoding the MexAB-OprM, MexCD-OprJ, MexEF-OprN, MexXY, and MexHI-OpmD drug efflux systems were kindly provided by Dr. Yuji Morita and Dr. Tomofusa Tsuchiya (21,24).

### Susceptibility test

MIC was measured using the broth microdilution method. We added 5 *μ*L of antimicrobials into 96-well plates with 95 µL of bacteria-containing medium and incubated them at 37°C for 18 hrs. Antimicrobials were prepared according to the Clinical and Laboratory Standards Institute (CLSI) (25). Bacteria were cultured in CAMHB or ID-CAMHB overnight at 37°C with shaking and finally tested at 5 × 10^4^ CFU/well after preparation to McFarland standard turbidity of 1 (± 0.1). For the combination of PA*β*N and cefiderocol, the checkerboard method (26) was used. Five µL each of cefiderocol and PA*β*N were added into a 96-well plate with 90 *μ*L of the bacteria-containing medium and incubated at 37°C for 18 hrs. Twofold dilutions of each compound were added into 96-well plates at various concentrations as indicated (Fig. 2). Bacteria were tested at 5 × 10^4^ CFU/well as MIC measurement.

### Growth curve measurement

Cultures of PAO1 and Δ*mexAB-oprM* were incubated overnight at 37°C with shaking, then diluted and added to CAMHB and ID-CAMHB to obtain an OD_600_ of 0.1. Then they were incubated with shaking at 37°C using Infinite M200 Pro (Tecan), and the absorbance in OD_600_ was measured over time.

### Chemicals

Cefiderocol was obtained from MedKoo Biosciences, Inc., (USA, Morrisville). Ceftazidime hydrate, cefepime hydrate, norfloxacin, Chloramphenicol, and PA*β*N were obtained from Sigma-Aldrich Co. LLC. (USA, St. Louis). Meropenem was obtained from LKT Labs, Inc. (USA, St. Paul). Imipenem was obtained from Banyu Pharmaceutical Co.Ltd. (Japan, Tokyo).

## Transparency declarations

None to declare.

